# Getting in shape and swimming: the role of cortical forces and membrane heterogeneity in eukaryotic cells

**DOI:** 10.1101/198523

**Authors:** Hao Wu, Marco Avila Ponce de León, Hans G. Othmer

**Affiliations:** School of Mathematics, 270A Vincent Hall, University of Minnesota.

**Author notes:** Supported in part by NSF Grants DMS 0817529 and 1311974, NIH Grant #54-CA-210190, the Newton Institute and the Simons Foundation.

**Keywords:** Low Reynolds number swimming, Self-propulsion, Membrane tension gradients Heterogeneous membrane, Boundary integral method

## Abstract

Recent research has shown that motile cells can adapt their mode of propulsion to the mechanical properties of the environment in which they find themselves – crawling in some environments while swimming in others. The latter can involve movement by blebbing or other cyclic shape changes, and both highly-simplified and more realistic models of these modes have been studied previously. Herein we study swimming that is driven by membrane tension gradients that arise from flows in the actin cortex underlying the membrane, and does not involve imposed cyclic shape changes. Such gradients can lead to a number of different characteristic cell shapes, and our first objective is to understand how different distributions of membrane tension influence the shape of cells in a quiescent fluid. We then analyze the effects of spatial variation in other membrane properties, and how they interact with tension gradients to determine the shape. We also study the effect of fluid-cell interactions and show how tension leads to cell movement, how the balance between tension gradients and a variable bending modulus determine the shape and direction of movement, and how the efficiency of movement depends on the properties of the fluid and the distribution of tension and bending modulus in the membrane.

*Dedicated to the memory of Karl P. Hadeler, a pioneer in the field of Mathematical Biology and a friend and mentor to many*.

## 1 Introduction

Movement of cells, either individually or collectively, plays an important role in numerous biological processes, including development, the immune response, wound healing, and cancer metastasis (Nürnberg *et al.* 2011). Single-cell organisms exhibit a variety of modes for translocation, including crawling, swimming, or drifting with a fluid flow. Some prokaryotes such as bacteria use flagella, while eukaryotes such as paramecia use cilia to swim, but both types can only use one mode. However other eukaryotes, such as tumor cells, are more flexible and can adopt the mode used to the environment in which they find themselves. For instance, whether a single-cell or collective mode of movement is used in tissues can depend on the density of the 3D extracellular matrix (ECM) in which cells find themselves (Haeger *et al.* 2015). This adaptability has significant implications for developing new treatment protocols for cancer and other diseases, for it implies that it is essential to understand the processes by which cells detect extracellular chemical and mechanical signals and transduce them into intracellular signals that lead to force generation, morphological changes and directed movement.

The two primary modes of eukaryotic cell movement on surfaces or in the ECM are called mesenchymal and amoeboid (*cf.* Figure 1.1) (Friedl & Wolf 2010, Binamé *et al.* 2010). The former is used by fibroblasts and tumor cells of epithelial origin, and typically involves strong adhesion to the substrate. In 2D movement is by extension of relatively-flat lamellipodia at the leading edge, whose protrusion is driven by actin polymerization. Growth of this structure is understood in terms of the dendritic network hypothesis, which posits that filaments are nucleated at the membrane and treadmill as in solution, and that the densely-branched structure of the network arises via nucleation of branches on existing filaments mediated by a protein called Arp2/3 (Pollard *et al*. 2000). Force is transmitted to the environment via integrin-mediated focal adhesions that are connected to the internal network (the cytoskeleton or CSK) via stress fibers. Movement frequently involves proteolysis of the ECM to create a pathway (Sanz-Moreno *et al.* 2008).

**Fig. 1.1.**
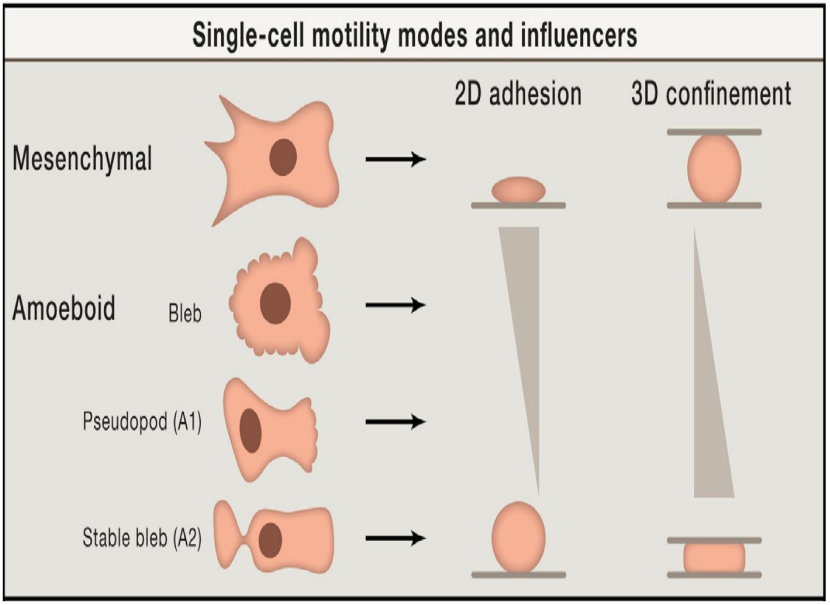
A summary of the different modes of movement in different environments and under different substrate properties. Modified from Welch (2015).

In contrast, the amoeboid mode utilizes a less-structured CSK that lacks stress fibers, and involves lower adhesion to the substrate. Proteolytic degradation of the ECM is generally not used, and cells adopt a more rounded cell shape, often with a highly contractile ‘tail’ called the uropod (Lämmermann *et al.* 2008). Two sub-types of amoeboid motion are known. In one, cells move by generating a rearward flow in the cortex – the cross-linked filamentous actin (F-actin) network that is linked to the membrane. As described later, the drag force created by the rearward flow leads to a reactive tension gradient in the membrane that propels the cell forward, and this is called the ‘tension-’ or ‘friction-driven’ mode. In another type, cells move by blebbing, in which cycles of extension of the front and retraction of the rear as shown in Figure 1.2(b) are used. How the spatial localization of blebs shown there is controlled is not understood, and some cells exhibit random blebs over their surface (*cf.* Figure 1.2(a)), which leads to no net translocation. Another variation of blebbing called ‘stable-bleb’ or ‘leader-bleb’ migration is used by certain embryonic cells that form a balloon-like protrusion at their leading edge (cf. Figure 1.1) and can move rapidly (Maiuri *et al.* 2015, Ruprecht *et al.* 2015, Logue *et al.* 2015).

**Fig. 1.2.**
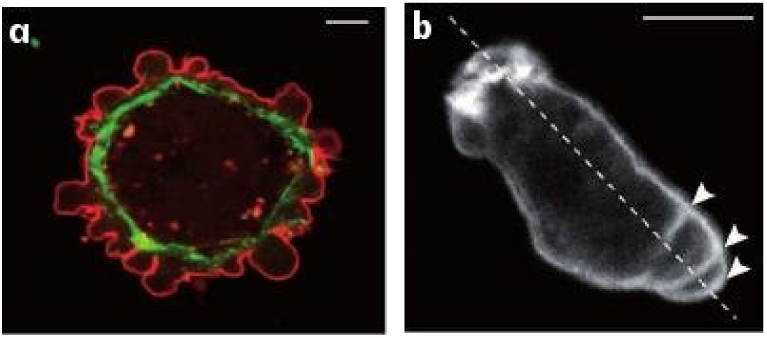
(a) Blebbing on a melanoma cell: myosin (green) localizes under the blebbing membrane (red) (b) The actin cortex of a *Dictyostelium discoideum* (Dd) cell migrating to the lower right. Arrowheads indicate the successive blebs and arcs of the actin cortex (Charras & Paluch 2008).

The amoeboid mode is widely used, and when the environment is less favorable to mesenchymal movement, due *e.g.*, to changes in the adhesiveness of the substrate, cells compensate by undergoing a ‘mesenchymal-to-amoeboid’ transition (MAT) (Friedl & Alexander 2011, van Zijl *et al.* 2011). Leukocytes can use the mesenchymal mode in the ECM, but can also migrate *in vivo* in the absence of integrins, using a ‘flowing and squeezing’ mechanism (Lämmermann *et al.* 2008). Dd moves in a cyclic AMP (cAMP) gradient either by extending pseudopodia or by blebbing, and cells monitor the stiffness of the surroundings to determine the mode: pseudopodia in a compliant medium and blebbing in stiffer media (Zatulovskiy *et al.* 2014). Finally, some cells move only by blebbing. Certain lines of carcinoma cells don’t move on 2D substrates, but in a confined environment they polarize spontaneously, form blebs, and move efficiently (Bergert *et al.* 2012).

A third, less-studied mode of movement is by unconfined swimming in a fluid. Dd cells and neutrophils both do this (Barry & Bretscher 2010), presumably to move through fluid-filled voids in their environment. A model of swimming by shape changes has been analyzed in (Wang & Othmer 2015a), where it is shown that Dd can swim by propagating protrusions axially. The model gives insights into how characteristics of the protrusions such as their height affect the swimmer’s speed and efficiency. Simplified models for movement by repetitive blebbing, using what can be described as a ‘push-pull’ mechanism, have been analyzed and the efficiency of movement determined (Wang & Othmer 2015b; 2016). However, it is also observed that Dd cells can swim without shape changes or blebbing for several body lengths (Howe *et al.* 2013), and the authors state that ‘Simply put, we do not understand how these cells swim, and therefore how they move.’ Herein we show how swimming can arise by maintaining an axial tension gradient in the membrane, and this would be difficult to detect at the macroscopic level.

Crawling and swimming are the extremes on a continuum of strategies, and the variety of modes used in different environments raises questions about how mechano-chemical sensing of the environment is used to control the evolution of the CSK (Renkawitz & Sixt 2010). Protrusions and other shape changes require forces that must be correctly orchestrated in space and time to produce net motion – those on cells in Figure 1.2 (a) are not, while those in (b) are – and to understand this orchestration one must couple the intracellular dynamics with the state of the surrounding fluid or ECM. Tension in the membrane and cortex has emerged as an important determinant in the orchestration, whether in the context of undirected cell movement, or in movement in response to environmental cues. Experimental observations on several modes of tension-driven movement are described in the next section.

## 2 The basis of tension-driven movement

To describe recent experimental results we introduce some terminology. We consider a cell as a three-layered structure comprised of the plasma membrane, the cortex, and the remainder (cytosol, nucleus and other components). The membrane is a lipid bilayer ~10 nm thick, and the cortex, which is 200-300 nm thick, is composed of an F-actin network cross-linked by filamin and bound to the motor protein myosin-II (myo-II), which can confer rigidity to the cortex, but can also contract and exert tension in the cortex. Membrane-bound proteins such as myo-I – a small motor protein that binds to both actin and the membrane (Dai *et al.* 1999) – or linker proteins such as ERMs (ezrin, radixin, and moesin), tether the cortex to the membrane and exert a normal force on the membrane. However the connection is dynamic and when the cortex flows it can slide tangentially under the membrane (Hochmuth *et al.* 1996, Dai & Sheetz 1999) and exert a tangential stress on the membrane. A detailed force balance done later shows how this can lead to swimming without either blebbing or shape changes.

Two recent papers describe a similar phenomenon – tension-driven motion of cells in confined environments – using human dermal fibroblast cells (Liu *et al.* 2015), or zebrafish germ-layer cells (Ruprecht *et al.* 2015). Liu *et al.* (2015) identify two morphologies, one type – A1 – has a rounded body and a small leading edge, and the other – A2 – has a more ellipsoidal body with a large uropod (Figure 1.2). They showed that slow mesenchymal cells undergo the MAT when the adhesion is low and the cells are confined between plates, and this leads to two distinct shapes and two types of fast migration. The first (A1) involves low contractility of the cortex and a local protrusion, and the second (A2), is a ‘stable-bleb’ type that involves high myo-II activity and involves a strong retrograde actin flow. Type A1 appears to require an external signal to polarize, whereas type A2 can appear spontaneously, as has been shown for other cell types as well (Lorentzen *et al.* 2011, Bergert *et al.* 2012). The authors suggest that the type A2 system may be bistable.

A different, permanently-polarized, stable-bleb shape can be obtained from a stable non-polarized blebbing cell by increasing the contractility in zebrafish progenitor cells (Ruprecht *et al.* 2015). The stable-bleb form involves cortical flow rates of 10’s of *µ*ms/min (Figure 2.1(a)), which would certainly induce an anterior-to-posterior cytoplasmic flow near the cortex, and thus a posterior-to-anterior flow in the center. The authors also postulate a high cortical growth rate at the front of a cell and a high disassembly rate at the rear (*cf.* Figure 2.1(b)). This hypothesis is supported by the fact that blebbistan (which inhibits myo-II contractility), latrunculin A (which leads to actin depolymerization), and jasplakinolide (which stabilizes actin networks) all inhibit polarization and cortical flow. The authors explain the transition from random blebbing to a stable-bleb shape as an instability of a spherical shape, which is adopted in the absence of surface contact, to fluctuations in the membrane. To date only a linear stability analysis of the problem has been done (Callan-Jones *et al.* 2016), and the results of that analysis are contrasted with the results herein in the Discussion section.

**Fig. 2.1.**
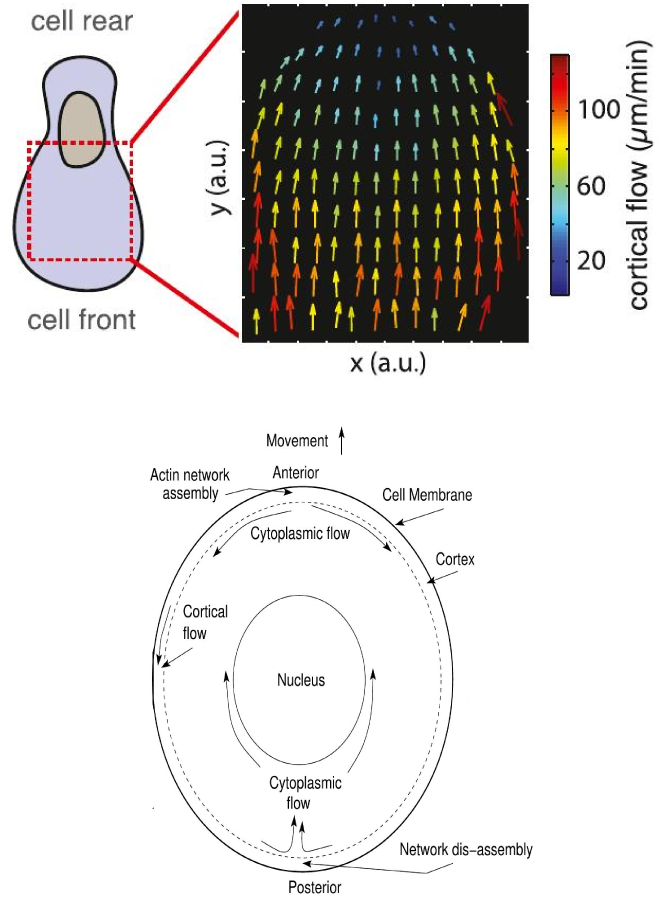
The measured cortical flow (top) (From Ruprecht, et al. Ruprecht *et al.* (2015)), and the postulated intracellular flows (bottom).

Thus there are two ‘stable-bleb’ cell morphologies in which a gradient of cortical density and myo-II contraction is used to generate a cortical flow and an axial pressure gradient (*cf.* Figure 2.2). How the flow is initiated, and in particular, whether it arises as an instability or requires contact with a substrate, is unknown. This mode of movement has only been demonstrated when cells are constrained to move between two horizontal barriers, and it has been suggested that substrate contact may open stretch-activated calcium channels and initiate modification of the cortex (Hung *et al.* 2016). Interestingly, some cells cannot move if they are only in contact with the substrate on the ventral (bottom) side, but will move when confined in a micro-channel (Bergert *et al.* 2012). This suggests that cortical flow may not arise when the cell is in contact with a surface on only one side, and this has been shown experimentally in Dd (Yumura *et al.* 2012).

**Fig. 2.2.**
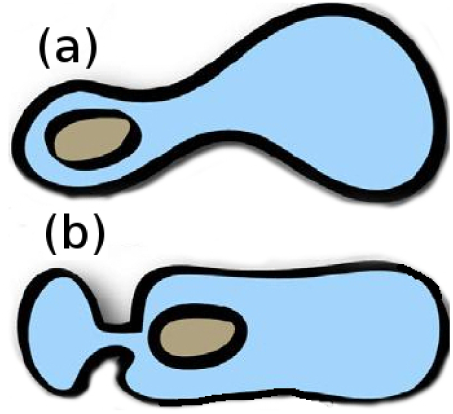
The two types of stable-bleb morphology in which movement in confinement is driven by tension gradients. (a) The Ruprecht-type, and (b) the Liutype A2.

To understand the origin of movement in these cells, recall that the cortex slips past the membrane, and in a numerous cell types, including Dd (Traynor & Kay 2007), leukocytes (Lee *et al.* 1990) and dendritic cells (Renkawitz *et al.* 2009), the membrane does not flow in a cell-fixed coordinate frame – it merely translocates with the cell body. Thus the membrane functions more like an elastic than a viscous material, and the drag force due to cortical flow creates an opposed tension gradient in the membrane. Back-to-front tension gradients of the order of 5*pN/µm*^2^ have been measured in axons and keratocytes (Lieber *et al.* 2015), but only on the dorsal membrane of cells in surface contact on the ventral membrane. We will show that tension gradients in the membrane can generate movement without shape changes, which we call ‘swimming’, of a cell submerged in a fluid. The plausibility of such motion is supported by the observed surface-tension-driven Marangoni propulsion of a fluid droplet on a viscous fluid (Lauga & Davis 2012). In fact, by cutting the cell in Figure 2.1 along the long axis, and opening it up and stretching the free ends to infinity, the membrane becomes an interface between the interior and exterior fluids, and one has the configuration of the classical Marangoni problem. Of course reality is more complex here, since the cortex as described above is a dynamic structure, thin at the front and thick at the rear,

While the primary experimental evidence of tension-driven movement involves cortical flows, we show that it is the existence of a tension gradient, whether or not it involves a cortical flow, that drives movement. Thus the results herein will be applicable to a larger class of cells than those used in the experiments described above.

## 3 The shape problem for cells

### 3.1 The free energy functional and the shape equations

The observed shapes described above, in particular the stable-bleb types shown in Figure 2.2, raise a number of questions. Since amoeboid cells have a less-structured CSK, the cell shape is primarily determined by the distribution of internal forces in the membrane and the forces in the cortex. Thus the first question is what distribution of forces in the membrane and the cortex is needed to produce the observed shapes? Secondly, since directed cell movement requires cell polarization and cortical flow, what balances in the cortex amongst cortical thickness, the level of myo-II for contraction, myo-I for attachment to the membrane, and other factors, are needed to produce the cortical flow? Finally, how do the properties of the micro-environment affect the speed of movement, and whether a cell switches from tension-driven movement to blebbing-driven movement?

A model that incorporates a detailed description of the cortical network growth, myo-II, and cortical-network interactions, combined with the transport of actin monomers and other components in the CSK, will be very complex, and to date the cortex has been described as an active gel in simplified treatments of cortical flow (Joanny & Prost 2009, Prost *et al.* 2015, Bergert *et al.* 2015). While this produces some insights, the biochemical details are embedded in an active component of the stress tensor for the gel, and thus the relative importance of the individual processes mentioned above cannot be investigated. We also do not attempt to develop a detailed model here, but rather, we use an alternate high-level description of the cortex to investigate how cortical forces and heterogeneity of membrane properties determine the shape of cells in quiescent fluids, and how these factors determine the shape and speed of swimming cells. Since the forces are simply specified, we can consider both the case in which there is a cortical flow that generates stresses, and the case in which the cortex is under stress but not flowing with this approach.

For this purpose we separate mechanics from the details of the cortical structure by prescribing a force distribution on the membrane that reflects what is believed to occur in the cortex *in vivo*. This allows us to vary the cortical forces directly, whether they arise from cortical flows or simply from static tension gradients in the cortex. We do this first for cells in a quiescent fluid, where there are no shear stresses due to the fluids, and secondly for swimming cells, in which the interior and exterior fluids flow freely. The three-fold aim of the latter step is to show that cells can swim when subject to cortical forces, to determine how much the shape of a moving cell differs from that of a stationary cell, and to show how the shape differences depend on cell and fluid properties.

The determination of the steady-state shapes of vesicles and red blood cells has been thoroughly studied, both in the absence of fluid motion (Seifert *et al.* 1991, Seifert 1997), and in the presence of fluid motion (Veerapaneni *et al.* 2009, Zhao *et al.* 2010, Li *et al.* 2014, Kaoui *et al.* 2009, Ghigliotti *et al.* 2010, Thiébaud & Misbah 2013). In the former the shapes are computed as minimizers of the free energy of the membrane, typically given by a Canham-Helfrich functional described below, and in the latter they represent shapes that lead to minimum dissipation in the flow. However, vesicles have no cortical layer and red blood cells have a very thin layer of spectrin – which contains no molecular motors – attached to the membrane. When there is a cortical flow and membrane-cortex tethers are actively formed and broken, there are dissipative processes involved and the membrane-cortex forces are not conservative. While one can use a virtual work argument to determine stationary shapes when there are non-conservative forces, we avoid this as follows. To find the stationary shapes under prescribed forces, we define a free-energy functional for the membrane, which we treat as an elastic medium since this conforms with experimental findings in Dd (Luo *et al.* 2013), we compute its first variation with respect to a deformation, which gives the membrane force, and to this we add the cortical forces directly.

The membrane has four modes of deformation: dilatation, shear, bending and torsion, but only the bending mode is treated in general and we follow this practice here. In addition to the bending energy, which to lowest order is proportional to the square of the local curvature of the membrane, there are contributions to the free energy corresponding to the work associated with area and volume changes when these are conserved.

We let Ω ⊂ *R*^3^ denote the volume occupied by the cell and let 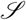 denote its boundary. We assume that 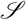 is a smooth, compact, two-dimensional manifold without boundary, parameterized by the map Φ: 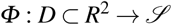, defined so that the position vector **x** to any point on the membrane is given by **x** = **x**(*u*^1^*, u*^2^) for a coordinate pair (*u*^1^*, u*^2^) ∈ *D*. We let **n** denote the outward normal on 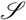, and define basis vectors on the surface via

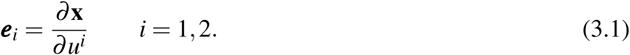

In general these are not normalized.

The free energy associated with bending, which was first set forth for membranes by Canham (1970) and later by Helfrich (1973), has the following form

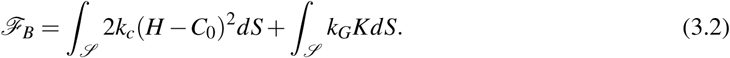

Here κ_1_ and κ_2_ are the principal curvatures, *H* = − (κ_1_ + κ_2_)/2 is the mean curvature, and *K* = κ_1_κ_2_ is the Gaussian curvature^1^. *C*_0_ is a phenomenological parameter called the spontaneous curvature, *k*_*c*_ the bending rigidity - which may be stress-dependent (Diz-Muñoz *et al.* 2016), and *k*_*G*_ the Gaussian rigidity, which may also vary over the membrane. When *k*_*G*_ is constant, the integral of the Gaussian curvature is constant if Ω does not change topological type under deformation, and the integral can be ignored.

Under the constraints of constant surface area *A*_0_ and volume *V*_0_ of the cell, the free energy takes the form

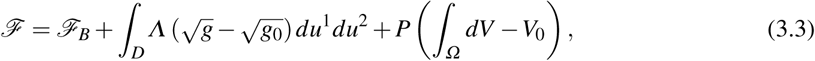

where *P ≡ p*_ext_ − *p*_in_ is the pressure difference across the membrane, which we assume is constant over the membrane. Typically *p*_in_ is a few hundred pascals higher than *p*_*ext*_ (Salbreux *et al.* 2012). The constant term *PV*_0_ simply translates the free energy and can be ignored, since it disappears after the first variation of (3.3) is taken. In the second-last integral over the boundary, *g* is the determinant of the metric tensor **g** of the surface, whose components are *g*_*i*__*j*_ = ***e**_*i*_ ⋅ **e***_*j*_, and *g*_0_ is its value in a reference or undeformed configuration in which the area is *A*_0_. The integral represents the energy needed to alter the area from it’s initial value, and *Λ* is the energy per unit area that arises when *A* is changed. Thus *Λ* has units of force/length, which defines a tension, but it is not a surface tension in the usual sense. Instead it is an in-plane stress of a two-dimensional surface, which can be seen as follows.

Consider a small x-y section of a thin flat plate of thickness h, and suppose there are no normal stresses to the section in the z-direction, which is orthogonal to the plate. Suppose the plate is an elastic material and let ***T*** be the stress tensor and let **ε** be the strain relative to a reference configuration. For small, uniform strains in the x-y plane on the four faces of the section one has

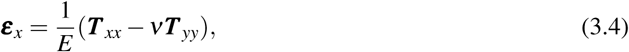

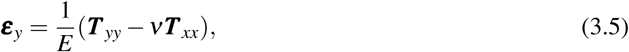

where *E* is Young’s modulus and ν is the Poisson ratio of the material. From these it follows that

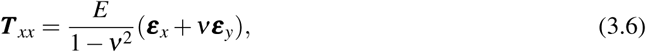

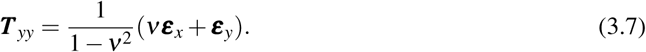

If the stress and strains in the x and y directions are equal and uniform in the z-direction, one can define a tension *T* as

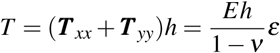

where ε = **ε**_*x*_ + **ε**_*y*_. Experimental results are usually reported in terms of a tension, but the foregoing shows that this assumes a local isotropy of the stress and strain. Evans & Needham (1987) define the tension as the average of the stresses along the principal directions of the surface, but these are rarely available. It is shown later that in the absence of imposed forces Λ is constant, and therefore the constant area term can be ignored as well.

In the absence of external forces, a stable equilibrium shape of a cell is a minimizer of 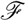, and thus a solution of 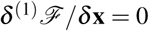 for any infinitesimal deformation **x** = **x**_0_ + ***ϕ*** + *ψ***n**, where 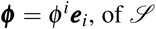. This leads to the following shape equations for the normal and tangential components of the membrane force.^2^

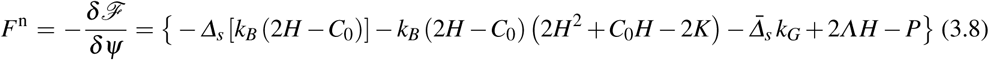

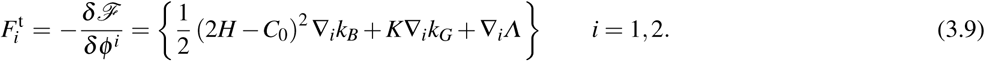

Here *∆*_*s*_ and 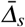 are two surface Laplacians and ∇_*i*_ is the component of the surface gradient, resp. In the first equation one sees that Λ enters the normal component via the term 2*ΛH*, which couples areal distension to the curvature in the normal component of the force. In light of how the variation is defined, the resultant forces are defined per unit area.

To simplify the equations we assume hereafter that *C*_0_ = 0, which reduces the foregoing to

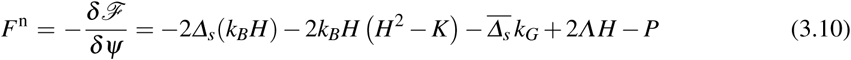

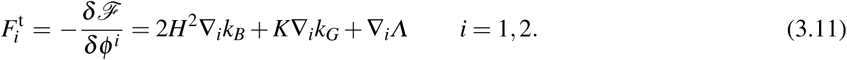

When the bending and Gaussian rigidities are constant these simplify to

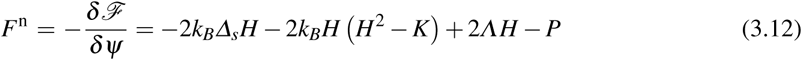

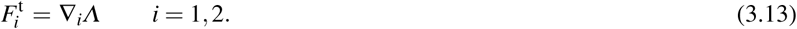

In any case the forces vanish at critical points of the energy, and stable shapes correspond to local minima of the free energy.

When written in (*u*^1^, *u*^2^) coordinates, the first term in *F*^n^ defines a fourth-order differential operator, and the critical points of the energy cannot in general be found analytically. Since the membrane is embedded in a quiescent viscous fluid, we define a fictitious relaxation process for the evolution of the shape from its initial values in which we suppose that the dominant force is a viscous drag, and we neglect inertial effects. This leads to the system

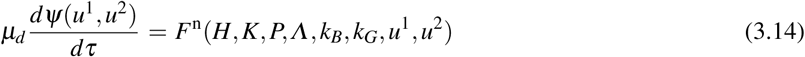

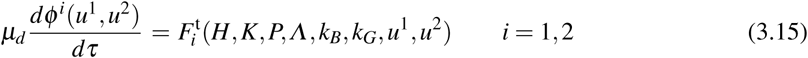

where *µ*_*d*_ can be thought of as a drag coefficient with the dimension of the viscosity per unit length.

### 3.2 The shape equations under cortical forces

To incorporate cortical forces, which are non-conservative, we add these to the intrinsic membrane forces as prescribed normal and tangential forces per unit area, the components of which are denoted *f*^n^ and 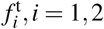. Thus the equations to be solved for a 3D shape are

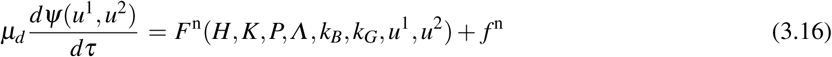

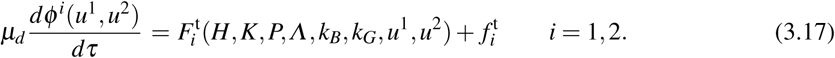

Since the normal force is directed inward and the normal vector is directed outward, *f*^n^ < 0. When the cortical forces are incorporated the resulting evolution is no longer a gradient flow, and one simply looks for steady states of (3.16) and (3.17), which in general are not minimizers of the membrane free energy.

The equations can be cast into non-dimensional form by defining the * variables 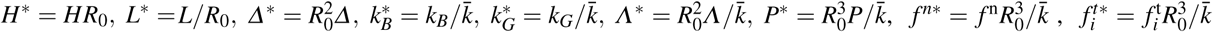 and τ∗ = τ/τ_m_, where *R*_0_ and 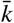 are the characteristic length and energy, resp., which we set equal to 1. Therefore, 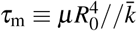 defines a characteristic time unit scaled by a constant characteristic bending rigidity unit 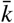. The resulting forms of (3.16) and (3.17) in unstarred variables are then

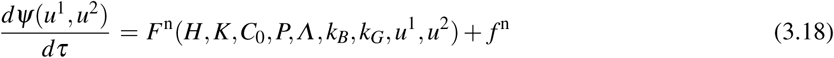

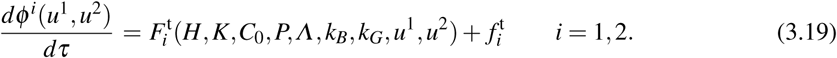

## 4 2D shapes

### 4.1 The evolution equations for 2D shapes

In order to understand the effects of membrane and cortical forces in determining the shape in the simplest possible context, we first perform an analysis of the cell shapes in two dimensions. In this case the domain is a 2D area, the boundary 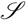 is a closed curve, and the area and volume constraints become perimeter and area constraints. The energy functional now reads

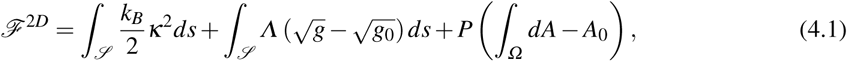

where κ is the curvature and *s* is the arc length on the boundary. The normal and tangential forces arising from the bending energy and constraints reduce to

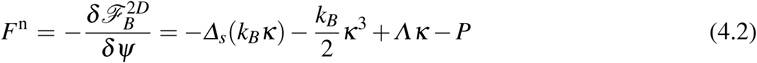

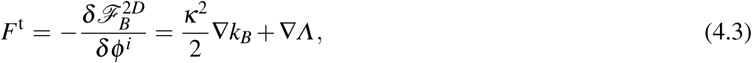

and the evolution equations (3.18) and (3.19) now read

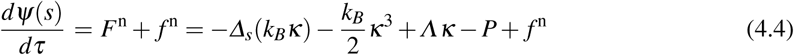

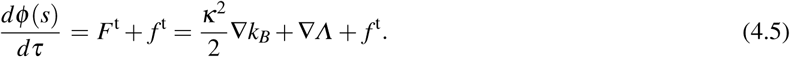

### 4.2 The numerical algorithm for solving the evolution equations

As a characteristic length scale of the problem we use an effective 2D cell size *R*_0_, which is defined via

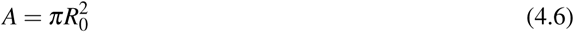

where *A* is the specified area enclosed by the cell contour, which has total fixed length *L*. The cell shape can then be characterized by the reduced area

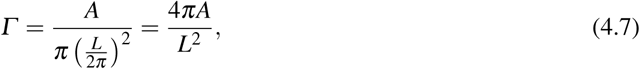

which is an intrinsic dimensionless parameter that expresses the ratio of the area of a given 2D shape to that of a circle of circumference *L* and a larger area.

To solve the 2D eqns (4.4) and (4.5), the boundary curve is discretized into *N* segments (Wu & Tu 2009) and we assume a ‘time’ step, τ_0_ and label the time sequence as τ_0_,2τ_0_, …, *j*τ_0_. The vectorial form of eqns (4.4) and (4.5) can be concisely transformed into

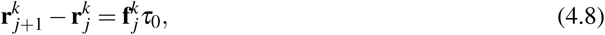

where 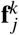 is the discrete form of the dimensionless force surface density. The geometric quantities in this equation can be discretized as:

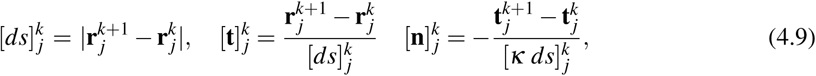

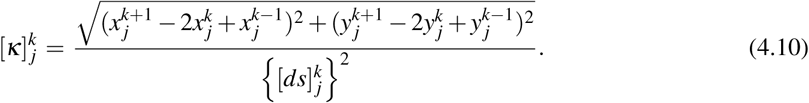

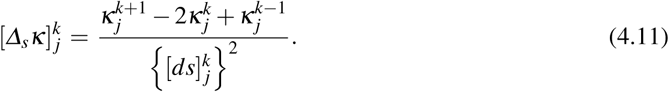

where 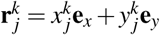 and 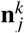 can also be obtained by rotating 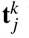 in 90° counterclockwise.

The local invariance of the boundary length is treated using a spring-like penalty method instead of introducing a spatially variable Lagrange multiplier, but our method is equivalent to it, as shown in Appendix B. One can re-define the areal energy density via an harmonic spring-like potential, which leads to a simple implementation. Let

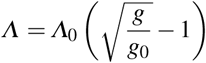

where *Λ*_0_ is a constant Lagrange multiplier. Then

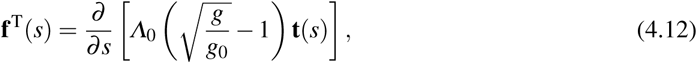

which can be rewritten as

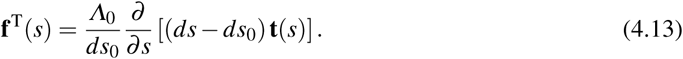

Eq(4.13) can be discretized as

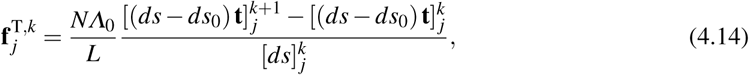

where *ds*_0_ = *L/N* and *L* is the total contour length of the 2D cell.

Because Λ_0_ can be considered as a membrane tension, one can define a new time scale τ_T_ = µ_o_*R*_0_/*Λ*_0_. This is to be compared to the membrane bending time scale τ_m_ defined earlier, and must be taken small enough in comparison to τ_m_ to ensure invariance of the boundary length on the membrane time scale. For most practical purposes τ_T_ = 10^−5^ − 10^−4^τ_m_ has proven to be sufficiently small (Wu *et al.* 2015; 2016, Ghigliotti *et al.* 2010, Thiébaud & Misbah 2013).

One could treat *Λ*_0_ as a true Lagrange multiplier, and develop a scheme to simultaneously solve for the shape and multipliers, but we simply fix a large *Λ*_0_ and check *en passant* that the arc length is conserved to the desired precision. The area constraint is treated as described in Appendix C. The stopping condition of the time-stepping calculation in (4.8) is 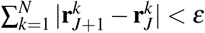 for a large enough integer *J* and a small enough number ε (ε = 10^−6^ is used here).

A summary of the computational procedure is as follows. First, the force distribution along the cell membrane is computed based on the current configuration of the cell. Second, the evolution equations (4.4) and (4.5) are used to obtain the new position in terms of the normal and tangential force distributions. Third, the first two steps are repeated until the stopping criterion has been satisfied, indicating that the steady state has been reached.

### 4.3 Computational results for 2D shapes

In this section we investigate the effects of spatial variations in the imposed cortical forces and in the bending modulus. Figure 4.1(a) shows representative shapes as a function of the magnitude of the dimensionless tether force 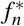 and the reduced area *Γ*, while Figure 4.1(b) shows the shapes as a function of the dimensionless tangential force 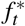 and *Γ*. In this figure and the following one, the property in question varies linearly from right to left of the shapes, with a minimum of zero at the left and the maximum given by the value on the x-axis. As a point of reference, if *Γ* = 1 all shapes are disks, irrespective of the variation of other properties. In both panels one sees that there is little effect on the shape when either force is less than 0.1, but a very strong effect for either force greater than 1. Shapes in the gray zone on the left are stable, but no stable shapes exist in the light yellow zone. Stable shapes are found for the entire range of the tangential force on the right.

**Fig. 4.1.**
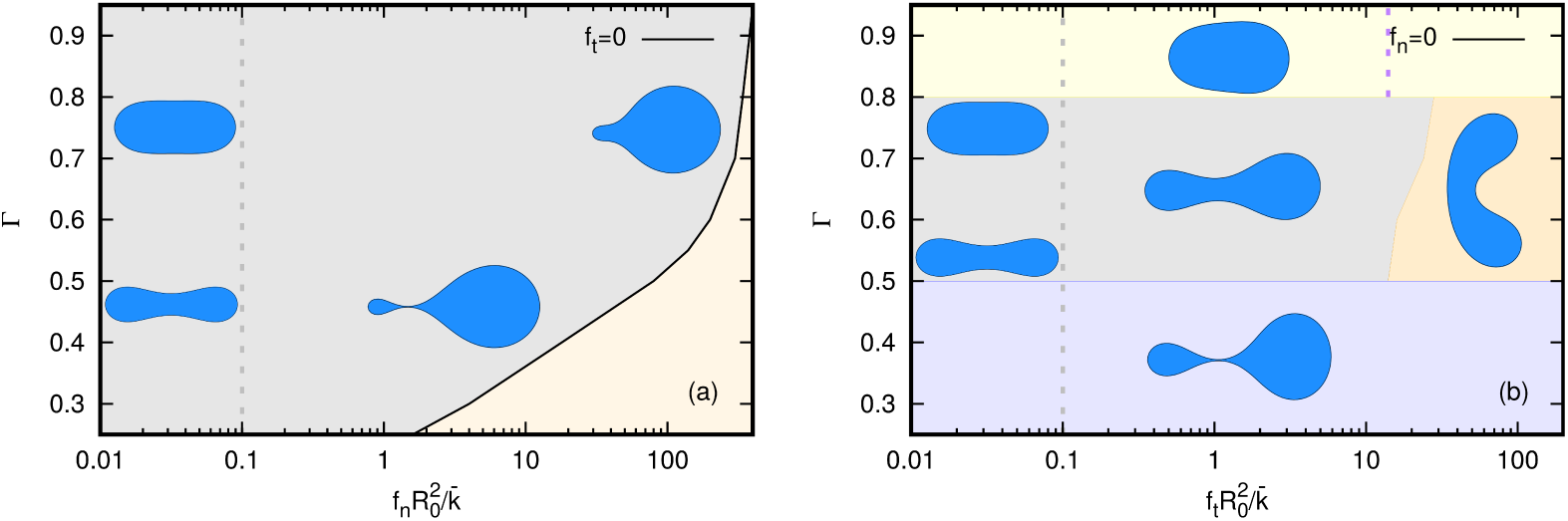
(a) A phase diagram showing cell shapes as a function of the dimensionless normal force 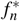 and the reduced area *Γ*. (b) A similar diagram for the dimensionless tangential force 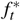 and the reduced area *Γ*.

The normal force (a) leads to pear-shaped cells at large *Γ* and shapes with two lobes connected by a narrow ‘bridge’ at *Γ* ~0.45, the latter somewhat similar to the Ruprecht ‘stable-bleb’ type shown in Figure 2.2(a). This shape only exists at a sufficiently small reduced area, when the shape under a small force is biconcave, and stems from the large normal force at the right end of the cell. A variety of other shapes can be obtained under different variations of the tether force. In particular, a symmetric two-lobed shape can be obtained by concentrating the normal force force at the center of the cell.

At 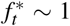 and small and intermediate *Γ* in the panel on the right, one sees that the large tangential component at the right end leads to a smaller lobe there, while the small tangential force at the left end leads to a larger lobe. These are again similar to the stable-bleb type observed experimentally, where it is suggested that the cortical tension is lowest at the front of the cell (the left end in the figure) and highest at the rear. Clearly the curvature is highest on the small lobe, which can be understood from the fact that a high tangential tension enters into the normal component of the membrane force equations. While the stable shapes are not necessarily minimizers of the area, the membrane-derived component of the forces drives the evolution toward a minimum. The tension energy is highest at the right end of the cell while the bending energy dominates the rear. Since the tension is lowest at the left end, the evolution there is driven by a tendency toward local energy minimization, which leads to larger radii to minimize the local curvature. If the magnitude of the tension increases further, the pear-like shape becomes unstable and evolves into a kidney shape with a shallow indentation or a kidney shape with a deep one, the latter shown in the light orange zone, depending on the applied magnitude and the reduced area. The kidney shape cannot be attained at low Γ due to the narrow neck that occurs around 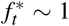, which precludes the shape changes involved in the transition to the kidney shape.

Experimental measurements of cortical tensions fall in a wide range, from 0.02 pN/*µ* for neutrophils to 4.1pN/*µ* for Dd (Winklbauer 2015). Estimates based on the number of linker proteins give a tether force of about 1 pN/*µ* (Diz-Muñoz *et al.* 2010). The results in Figure 4.1 are given in dimensionless terms, and a comparison of them with the experimental results requires the bending rigidity of a curve. Since the bending rigidity of a bar scales differently than that of a plate, one cannot use the latter – which is experimentally measured – to estimate the former. Thus a comparison of the model predictions with experimental results must await 3D computations of the shapes as a function of the applied forces.

To investigate the effect of a variable bending modulus, we vary it axially using the hyperbolic tanh function

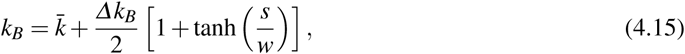

where *s* is the local length measured from the right end of the cell, 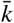 is the characteristic bending energy unit, ∆*k*_*B*_ is the bending rigidity difference between the right and left ends of the cell, and *w* is the width of the transition zone. In a thin-shell description of a material the Young’s modulus varies as the thickness cubed, and hence the axial variation imposed here could arise from the axial variation of the cortical thickness. It might also reflect a distribution of adsorbed or transmembrane proteins, and of glycolipids (Wu *et al.* 2013). Figure 4.2 shows that in a quiescent fluid a cell undergoes a shape transition from the biconcave shape to the pear shape induced by the variable bending rigidity. Since there are no imposed forces here the shapes are minimizers of the free energy. One sees that the cell ‘expands’ in regions of large rigidity since a region of higher *k*_*B*_ offsets the lower curvature in the free energy 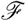.

**Fig. 4.2.**
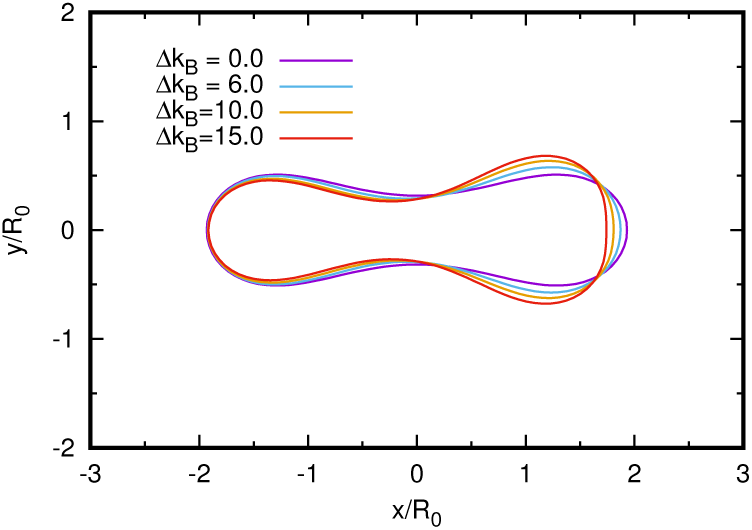
The dependence of shape on the magnitude of the variation in *k*_*B*_. The shape changes from biconcave to the pear shape as the axial variable bending rigidity increases. *Γ* = 0.6

## 5 Tension-driven swimming at low Reynolds number

### 5.1 The boundary integral equation

When the cell is submerged in a fluid and free to move, it generates extra- and intracellular flows, and here we assume that both the extra- and intracellular fluids are Newtonian, of density *ρ* and viscosity *µ*. In reality both fluids are undoubtedly more complex, but to demonstrate that movement ensues under tension gradients in the membrane it suffices to consider Newtonian fluids.

The Navier-Stokes equations for the intra- and extracellular fluid velocities **u** are (Childress 1981)

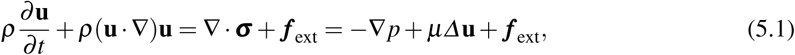

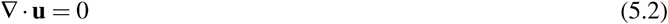

where

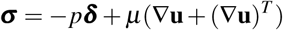

is the Cauchy stress tensor and **f**_ext_ is the external force field. We assume that the intra- and extracellular densities are equal, and thus the cell is neutrally buoyant, but allow for different viscosities in the two fluids. We further assume that there are no additional body forces and thus set *f*_ext_ = 0.

When converted to dimensionless form and the symbols re-defined, these equations read

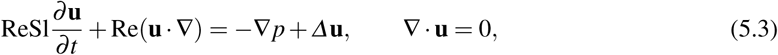

where the Reynolds number Re based on a characteristic length scale *L* and a characterisic speed scale *U* is Re = ρLU /*µ*. In addition, Sl = ω*L/U* is the Strouhal number and ω is a characteristic rate at which the shape changes. When Re ≪ 1 the convective momentum term in equation (5.3) can be neglected, but the time variation requires that ReSl = ωρ*L*^2^/*µ* < < 1, which implies that the initial shape changes must be slow enough. When both terms are neglected, which we assume throughout, a low Reynolds number (LRN) flow is governed by the Stokes equations, now in dimensional form,

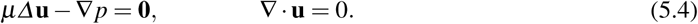

Here we consider small cells such as Dd, whose small size and low speeds lead to LRN flows (Wang & Othmer 2015a), and in this regime time does not appear explicitly, there are no inertial effects, and bodies move by exploiting the viscous resistance of the fluid. In the absence of external forces on the fluid and acceleration of the swimmer, there is no net force or torque on a self-propelled swimmer in the Stokes regime, and therefore movement is a purely geometric process.

The solution of the Stokes equations via the boundary integral method is a well-studied problem for red blood cells or vesicles in a pressure-driven flow (Zhao *et al.* 2010, Veerapaneni *et al.* 2011, Rahimian *et al.* 2015), but here the tension that arises from the active contraction of the cortex drives the flow. In the boundary integral method (BIM) one uses an Eulerian description for the velocity field of the fluid and a Lagrangian description for the configuration of the membrane. We assume that the interior and exterior viscosities are equal and that the velocity at infinity vanishes, and therefore the solution has the representation (Pozrikidis 1992; 2003)

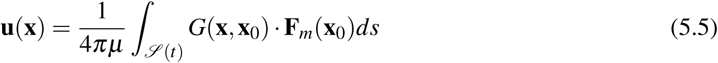

where

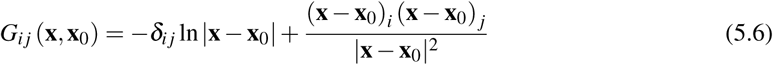

is the Green’s function in two space dimensions and **F**_*m*_ is the force exerted by the membrane on the fluid. We assume continuity of the velocities across the membrane, and mechanical equilibrium at the membrane, and therefore the force balance reads

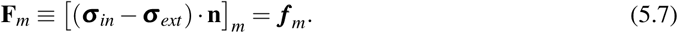

The right-hand side of (5.7) is the force on the membrane given by eqns (4.4) and (4.5), and thus one has to solve the integral equation, the shape equation, and the tangential force balance to determine the interior, exterior and boundary velocity fields.

When 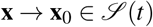, the Green’s function in (5.6) diverges, but it is a weak singularity, and can be removed by using the singularity subtraction transformation (SST) (Farutin *et al.* 2014), which is based on the integral identities

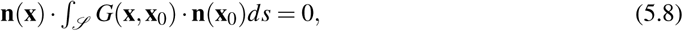

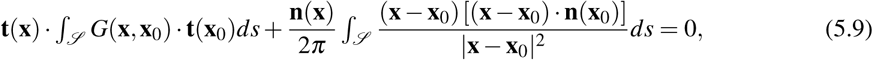

where **t** is the tangent vector to the contour 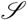. The first identity follows from the divergence theorem and the second from Stokes’ theorem. The SST technique for the single-layer kernel *G*(**x**, **x**_0_) can be applied as follows. Rewrite the integral in (5.6) as

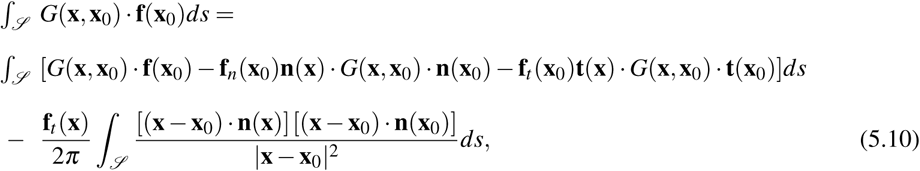

where **f**(**x**_0_) = *f*_*n*_(**x**_0_)**n**(**x**_0_)+ *f*_*t*_ (**x**_0_)**t**(**x**_0_) and **f**_*n*_(**x**_0_) = (**f**(**x**_0_) ⋅ **n**(**x**_0_))**n**(**x**_0_) and **f**_*t*_ (**x**_0_) = (**f**(**x**_0_) ⋅ **t**(**x**_0_))**t**(**x**_0_) are the normal and tangential parts of the force **f**(**x**_0_). Both integrals on the RHS of (5.10) have no singularities, that is they both go to zero when **x** approaches **x**_0_ along the contour 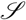. The second term in (5.10) tends to zero when **x**_0_ approaches **x** along the contour because 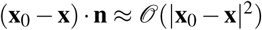.

### 5.2 The numerical algorithm for cell swimming

Step 1: Following the dimensionless quantities denoted with a star symbol in Section 3.2, the dimensionless surface force in unstarred variables **f**(**x**_0_) can be expressed in the component as the right hand side (RHS) of eqns (4.4) and (4.5) in terms of the current configuration of the cell.

Step 2: By substituting the instantaneous surface force distribution into the dimensionless boundary integral in unstarred variables along the cell shape contour, we can obtain the velocity of each material node on the membrane

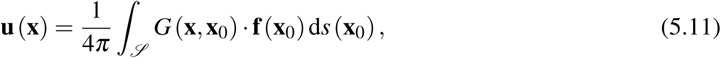

Step 3: The motion of cell membrane is obtained by solving the dimensionless time evolution equation in unstarred variables

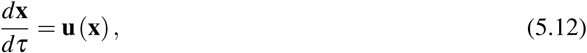

for each material node **x** lying on the membrane and the new cell configration is updated at each time step using an explicit Euler scheme

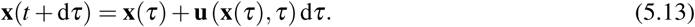

Step 4: Repeat the former three steps until the stopping criterion has been satisfied, indicating that the steady shape and swimming velocity have been obtained.

The discretization procedure and validation of the method can be found in Appendix D.

### 5.3 Computational results for cell swimming

To compute the velocity field in the interior and exterior fluids we introduce a square Eulerian grid with a specified degree of refinement – the mesh size can be taken significantly smaller than d*s* according to the accuracy required. Since the Green’s function is singular when the target point coincides with the source point, a small strip (of order d*s* around the boundary is excluded from computation of the velocity field. The Eulerian lattice grid points do not coincide with the Lagrangian nodes on the membrane in general, and we need only to evaluate the distance between the source point (lying on the membrane) and the target point (lying on the square lattice grid). For each point **x**, the velocity field is evaluated by using (5.11), where the integral along the membrane is performed exactly. A simple but accurate algorithm to judge the cellular interior and exterior is given in Appendix E.

Figure 5.1 shows the velocity field outside the swimming cell in a cell-fixed frame. The interior flow (not shown) indicates that tension driven swimming can form a microcirculation flow inside the cell, which is consistent with the biological conjecture shown in Figure 2.1. Figure 5.2(a) shows that the swimming velocity as a function of the applied forces is linear for all normal forces (a), and linear for small tangential forces (b). For large tangential forces, the speed drops abruptly in the transition to a kidney shape. The plateau shown in (b) is not precisely flat, but the values are close.

**Fig. 5.1.**
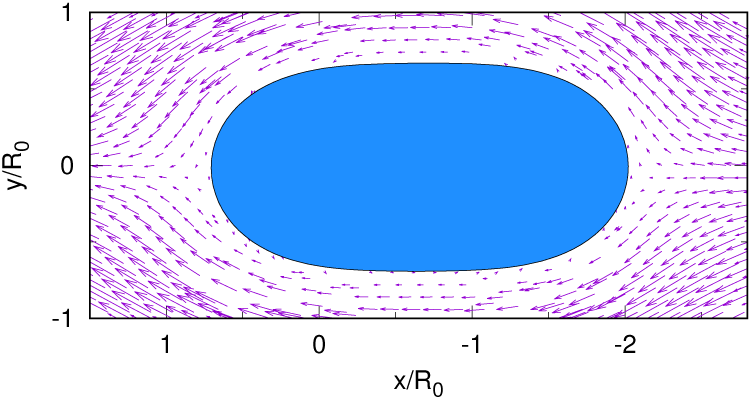
The velocity field outside the swimming cell.

**Fig. 5.2.**
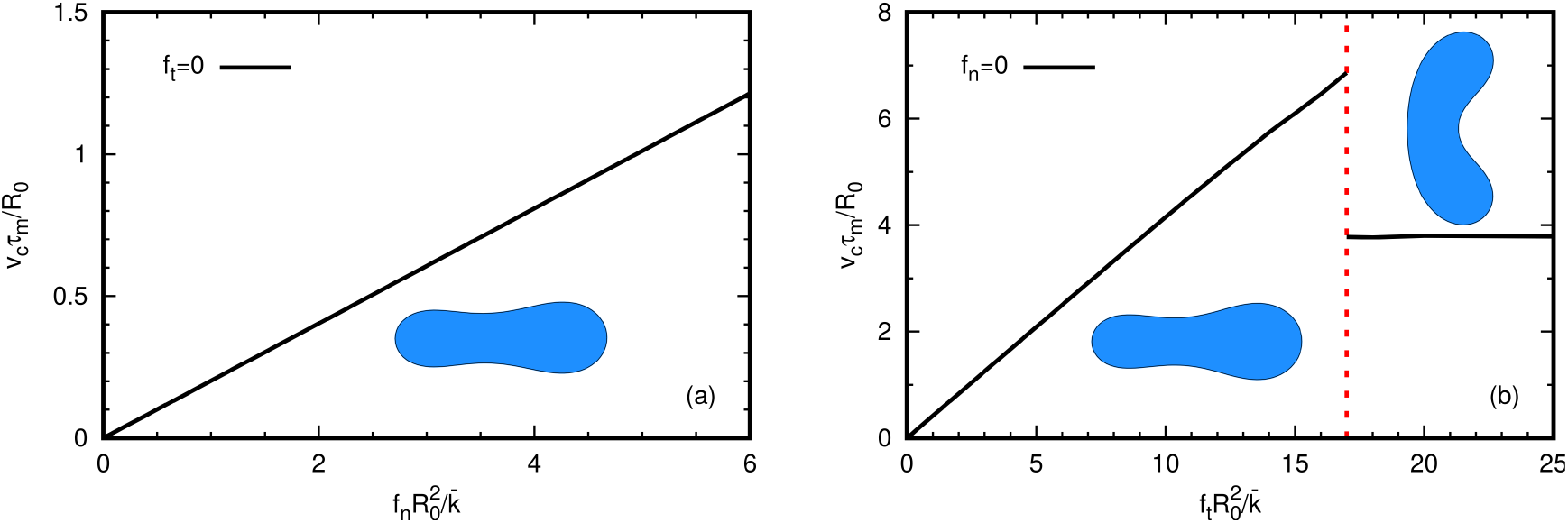
The swimming velocity as a function of the applied forces. The gradient of the forces is reversed from earlier figures, and the cells move to the right. In the both diagrams the cell has a reduced area *Γ* = 0.6.

Figure 5.3 shows the velocity and evolution of cell shape as a function of time. Initially the shape is biconcave, but evolves to the pear shape induced by the variable bending rigidity, and the corresponding swimming velocity changes with time. As in Figure 4.2, the lobe size depends on the bending rigidity difference between the front and the rear of the cell – the larger the difference, the larger the difference in lobe size. The inset shows the force distribution on the boundary, where one sees that stress concentrates on the large lobe.

**Fig. 5.3.**
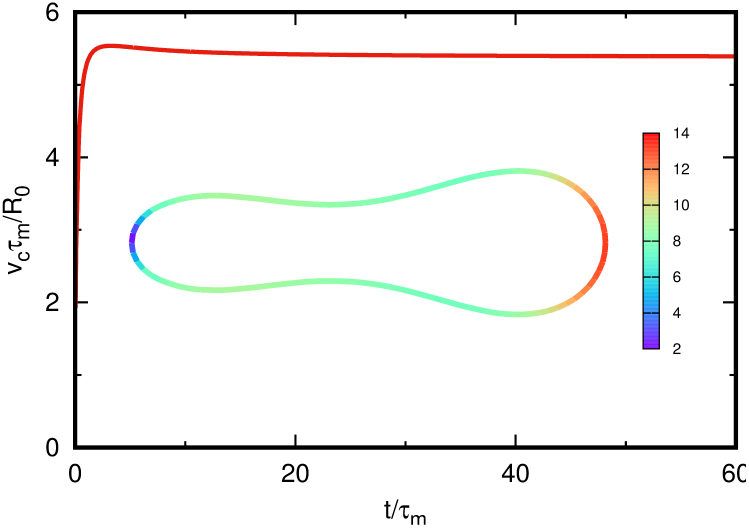
The velocity, shape and surface force distribution on the cell. The cell shape has a reduced area *Γ* = 0.5 with *f*_*t*_ = 0.0, *f*_*n*_ = 0.0, and ∆*k*_*B*_ = 6.0. The cell moves in the direction of the large lobe.

## 6 Discussion

Metastasis of a cancerous growth, which involves development of secondary malignant growth distinct from the primary tumor, accounts for the poor prognosis in many cancer types, and thus understanding how cells move in various environments is an important step toward devising medical treatment strategies for inhibiting metastasis. Because many cell types can use different modes ranging from crawling to swimming, it is important to understand how the micro-environment a cell finds itself in influences the mode it chooses. The shape a cell adopts in different environments is dependent on both intra- and extracellular forces on the membrane, and the balance between them can in turn determine the mode of movement via feedback of mechanical forces on the structure of the cytoskeleton and cortex. Earlier we described how certain cell types generate cortical flows that can produce motion, but there is little understanding of the quantitative relationships between the various forces, cell shape and movement.

Recently Callan-Jones *et al.* (2016) addressed the question of how cortical flows affect the shape of cells analytically, via a perturbation analysis of a spherical cell. They find that for small perturbations of a sphere the cortical force is the primary determinant of the shape, and showed that the flow can lead to axial asymmetries of the shape, similar to those observed. However a complete treatment of the control of cell shape and movement is currently beyond reach, either analytically or computationally, and in this paper we have addressed several simpler questions, which are (i) how do the shapes adopted by cells constrained in a quiescent fluid depend on prescribed cortical forces, and (ii) how does a cortical tension gradient affect the shape and speed of movement when it is unconstrained.

A primary objective of this work was to demonstrate that cells could swim without any shape changes, simply by using tension gradients in the membrane. To show this, we developed a two dimensional mechanical theory that captures the necessary cortex-membrane and fluid-membrane interactions. This enabled us to analyze the range of steady state shapes that could be adopted by membranes under tension distributions and whether these distributions could lead to swimming via an interaction with the fluid. We found that the resulting shapes closely resemble some of the shapes observed experimentally. Several new features of cell deformation and microswimming have emerged, including the following. (1) The pear shapes can be replicated by both the normal and tangential applied tension gradients, as well as by a variable bending rigidity. As the force or modulus variation is increased, the imposed asymmetry produces the transition from the non-polar bi-axial-symmetry (biconcave shape) to the polar uniaxial symmetry (pear shape or kidney shape). (2) The cell swimming velocity is linear as a function of either the normal or tangential forces, but the strong tension gradient in the tangential direction can further lead to a kidney shapes and cause the swimming velocity to abruptly decrease. (3) The velocity field within the cell forms two circulating loops which provide a mechanism for shuttling actin monomer from the rear to the front, as has been postulated by experimentalists, and is shown schematically in Figure 2.1.

Future work on 2D cells using the current model will focus on the interactions between the different factors with a view toward more accurately replicating the observed shapes and further investigating fluid effects such as differences in the interior and exterior viscositites. We believe that we will be able to obtain most of the shapes that are observed with proper modification of the parameters in the model. For example, some shapes probably arise via highly localized force distributions, and can probably be obtained by suitable modification of the applied tension distributions. Another aspect for study concerns movement in confined spaces, which can be done with the present methods if the boundaries are not too close to the cell, but which will require new methods when the cell is in direct contact with the substrate.

Although the present study provides a first step for the modeling of non-adhesive cell movement, several other questions deserve future consideration in order to capture a more realistic picture of tension driven swimming. For example,in the present work, we assumed a single layer integrating both effects of membrane and cortical layers whereas real cells are endowed with one lipid membrane layer and a separate cortical layer, and a nucleus that makes deformations more difficult. It will be of great importance to include the two separated layers and celluar nucleus in modeling for better comparison with experimental systems. Another future issue is to study the environment effects on cell migration, for example, what is the spatial confinement effect on cell swimming velocities and trajectories and what is the cell dynamics in a viscoelastic fluid. The last issue is that in nature cells migrate in a 3D environment, thus a more realistic model would incorporate the effect of the second principal curvature on the migration phenomena of interest in this paper. A 3D model would also allow for a more accurate description of the cortex-membrane and fluid-membrane interactions. For example, applying an anisotropic force distribution could lead to more realistic steady state shapes. A complete investigation of this effect will be reported in future work. Some preliminary results indicate that the cross section along the longest axis of the 3D shapes adopted by cell membranes under tension coincide with the two dimensional results shown here. Figure 6.1 below shows the equilibrium shapes of a membrane with and without an external tension gradient.

**Fig. 6.1.**
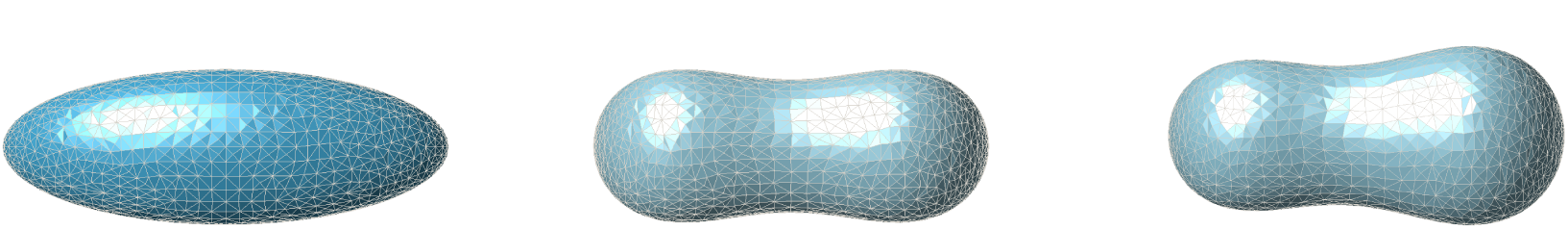
Here we exhibit the equilibrium shapes adopted by a cell membrane assumed to have an initial shape of a 3x1x1 ellipsoid shown at the left. The figure in the center is the typical biconcave shape adopted by a membrane in the absence of cortical forces. At the right we can see the asymmetric final shape that is adopted when the membrane is under a linear tension gradient, going from 10pN/*µm*^2^ in the back to 0 pN/*µm*^2^ in the front. The numerical method used to produce these shapes is based heavily on that of (Bonito *et al.* 2010). The computations were performed using the finite element software FELICITY due to Walker (2017).

## Acknowledgements

We thank Prof. Chaouqi Misbah for generously providing a basic simulation code written by Dr. Marine Thiébaud and originally used for the simulation of vesicle dynamics in an infinite channel (Thiébaud & Misbah 2013).

## Appendices

## A An outline of the derivation of the shape equation

In 1989 Ou-Yang & Helfrich (1989) computed the first, second and third order variations of the Helfrich energy under normal deformations of the membrane by using traditional tensor analysis. Later Capovilla *et al.* (Capovilla *et al.* 2003) applied the covariant geometry and Tu & Ou-Yang (2004) applied Cartan’s moving frame method to recalculate the variation of the original Helfrich free energy in both normal and tangential directions, respectively. Here, we extend these calculations to include a heterogeneous bending and Gaussian modulus.

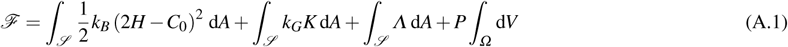

Note that the Gauss-Bonnet theorem is only valid for a constant Gaussian modulus, *k*_*G*_. If *k*_*G*_ is variable, then one cannot ignore the second term in the energy when performing the first order variation.

For the derivation we introduce the following symbols and definitions. The unit normal vector **n** points outward. *g* is the surface metric tensor, and *L* is the extrinsic curvature tensor. 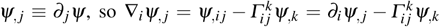 where 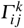 is the Christoffel symbol of the second kind, which is symmetric. *H* = −(κ_1_ + κ_2_)/2 is the mean curvature, *K* = κ_1_ κ_2_ is the Gaussian curvature, where κ_1_ and κ_2_ are the principal curvatures. The surface Laplacian is 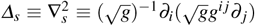 and 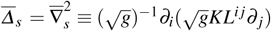 and *i, j* = 1, 2, throughout.

We will use the following identities of first order variations taken in both the normal and tangential directions,

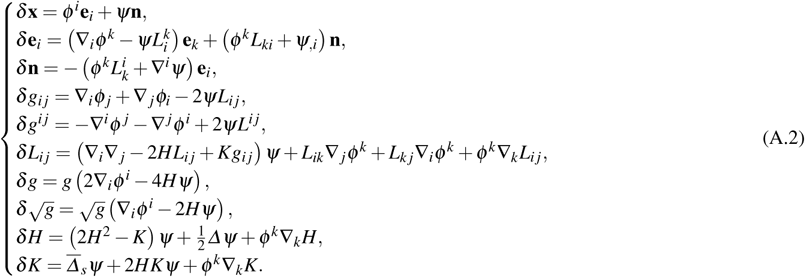

We can calculate the general variation of the modified Helfrich free energy as follows:

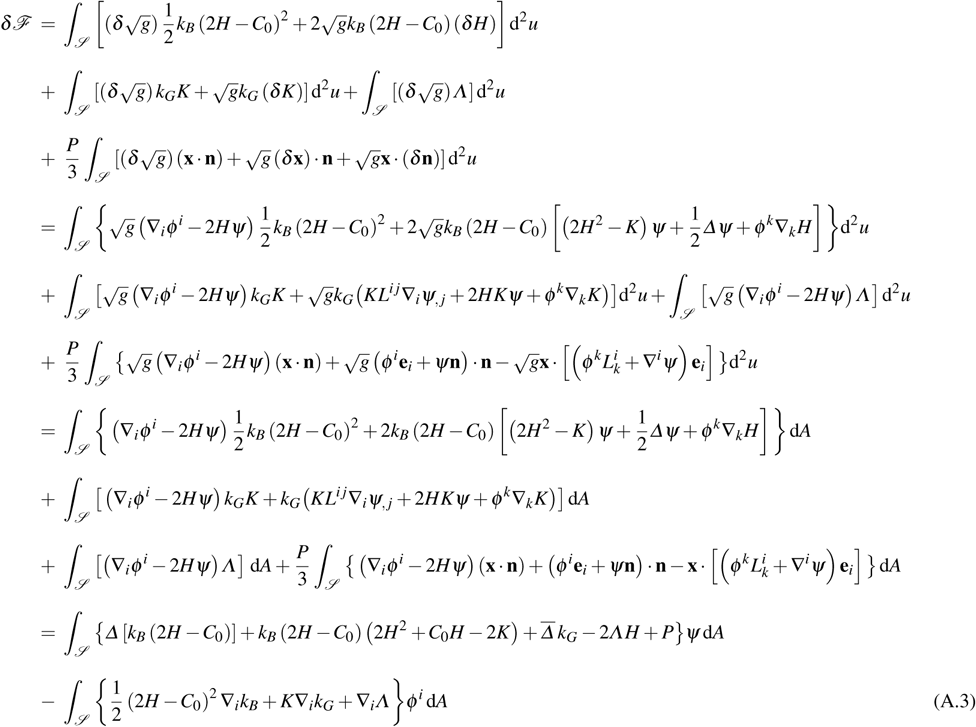

Thus, the two first functional derivatives (force components) in the normal and tangential directions are as follows:

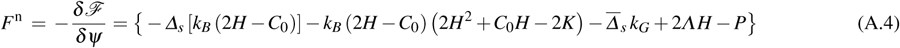

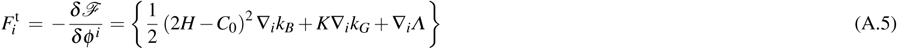

### B Area and arc length conservation via an harmonic potential

There are several ways in which the area conservation term in the free energy (3.3) can be written, and here we show one which has a simple interpretation in terms of springs between nodes and is equivalent to the form given in (3.3) under an appropriate definition of *Λ*. This method has been used by numerous authors (Henriksen & Ipsen 2004, Finken *et al.* 2008, Freund 2007, Thiébaud & Misbah 2013, Wu *et al.* 2015; 2016), and previous implementations of it have shown that the precision of this method depends on the choice of the coefficient *Λ*^*s*^(*u, v*). The area or arc-length can be conserved as precisely as desired by making *Λ*^*s*^(*u, v*) large enough, with the disadvantage that this may require taking a small time step in the numerical algorithm.

The method is based on the following formulation of the area constraint,

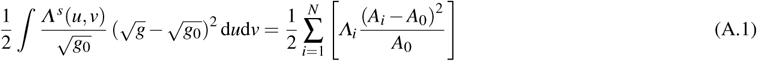

where *Λ*^*s*^(*u, v*) is fixed in the original reference frame. *g* is the metric on the deformed surface and *g*_0_ is the metric fixed in the original surface.

The equivalence between the spring-like penalty method and the Lagrange multiplier method is as follows. The variation of the areal energy under a perturbation can be written as

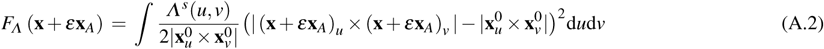

Therefore the first variation can be written

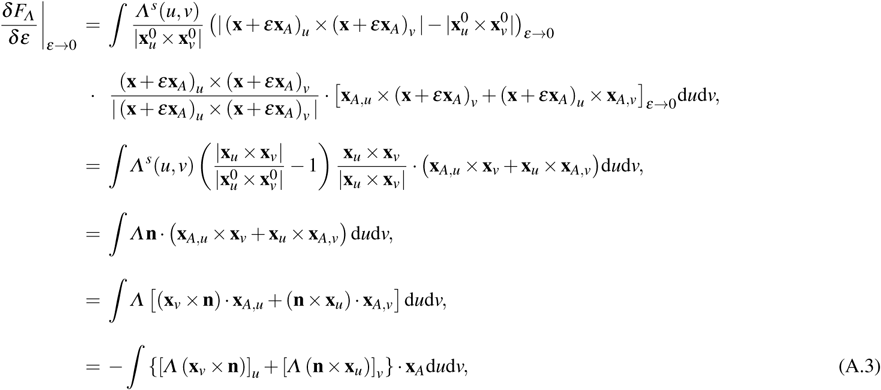

where

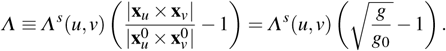

By using the vector product formula ***a*** × (***b*** × ***c***) = (***a*** ⋅ ***c***) ***b*** − (***a*** ⋅ ***b***) ***c***, we obtain

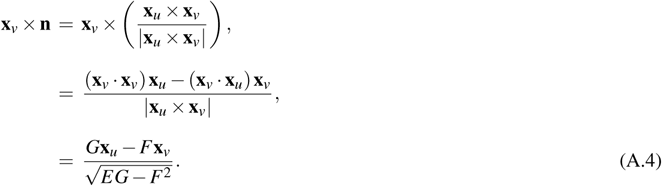

Similarly,

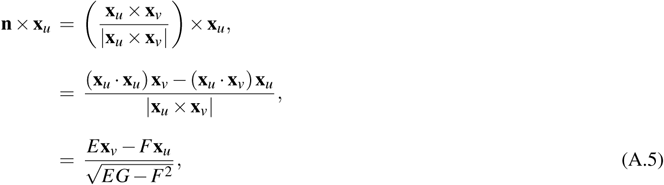

where *E*, *F* and *G* are the coefficients of the first fundamental form.

The surface divergence of a vector field is

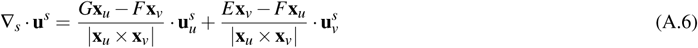

(do Carmo 1976), and the surface gradient of a scalar field is

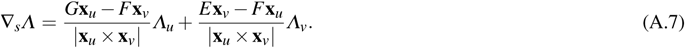

From the Weigarten equation, 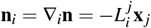, we have that, 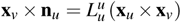 and 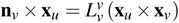, where we are assuming the Einstein summation convention. Because the mean curvature 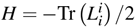, we can further obtain

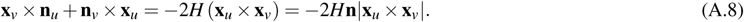

From equation (A.7) and equation (A.8), we easily have

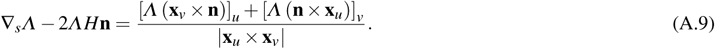

Therefore, equation (A.3) can be further calculated as

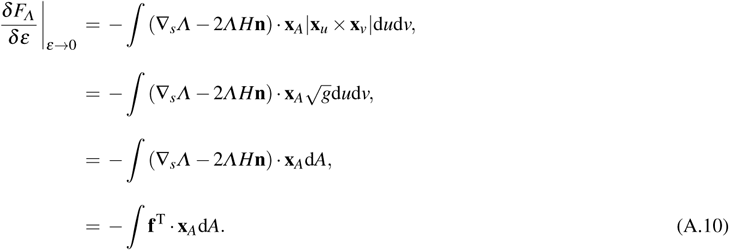

Thus, the spring-like penalty method is equivalent to the Lagrange multiplier method for the appropriate definition of *Λ*.

### C A global area or volume conservation condition

There is no Lagrange multiplier needed for the volume conservation in an incompressible fluid – the volume is automatically conserved if the divergence-free condition is met. However, in practice there is still some deflation in the process of numerical calculation. Thus a volume correction must be executed at each step, in order to precisely conserve the initial volume.

The traditional formula, which consists of summing all the volumes of the element triangular prisms has an obvious disadvantage in that it fails to calculate the volume when the center of mass is sometimes located outside the cell body. To avoid this disadvantage, we introduce the Minkowski identity for the volume of an N-dimensional geometric object in the space *R*^*N*^

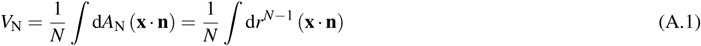

where *N* is the dimension of the geometric object. *V*_N_ and *A*_N_ are the volume and the area of the N-dimensional object.

To prevent the accumulation of errors of the area in 2D, or the volume in 3D, we implement a correction step due to (Freund 2007). We state this correction for the volume case, the area version is given in (Freund 2007). Suppose we apply a small perturbation **p** to the membrane surface. The volume can be precisely conserved as follows. Consider the functional

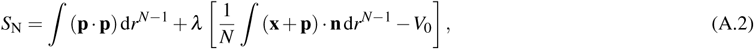

where *N* is the dimension of the geometric object and λ is a Lagrange multiplier to keep the volume conserved. A shape which minimizes the action above will satisfy that

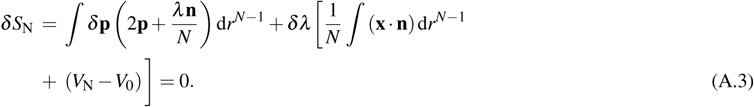

By solving equation (A.3), we obtain

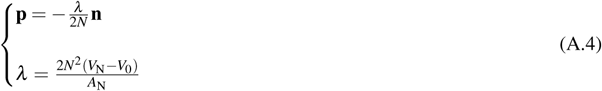

Thus, the corrected position vector to preserve the volume can be expressed as

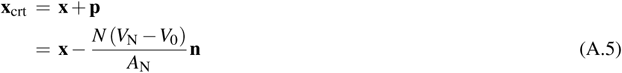

Numerical checks show that that this method can preserve the volume to within 10^−3^*V*_0_.

### D Discretization and parametrization implementation of the boundary integral method

The cell boundary curve is discretized into *N* segments with *N* points (Thiébaud & Misbah 2013, Wu *et al.* 2015; 2016). We label the *N* points in the boundary as 1, 2, … *, k* − 1*,k,k*+1, … *, N*. The discrete form of surface force 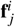 (bending part and tension part) can be expressed using the procedures in Section 4.2.

The velocity of each node on the cell membrane has the discrete form of the integral equation (5.11) in terms of the trapezoid rule and can be approximated using a finite difference scheme

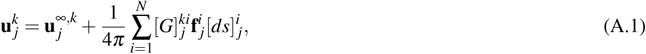

where 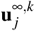 is a constant velocity in the *k*-th node at the *j*-th timestep and the bending part of 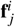 can be discretized using equation (4.10). The tension part of 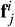 has been discretized in Section 4.2. The standard discretization of the free space Green’s function 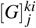 can be found in the book (Pozrikidis 2002).

Assuming the step of evolution time is τ_0_, the time sequence as τ_0_,2τ_0_*,…,* (*j* − 1)τ_0_*, j*τ_0_, (*j*+1)τ_0_*,…*. Equation (5.13) is transformed into

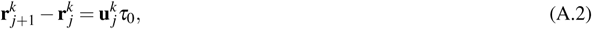

Finally, the new surface force on each node is estimated again from the new cell configuration 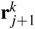, and the whole scheme above is repeated for a long time until a the stationary state is reached.

In this paper, the cell membrane contour is discretized using *N*_m_ = 120 nodes whose positions are updated at each time step of τ_0_ = 10^−4^τ_m_. The relative errors corresponding to the area, the perimeter and the reduced area are around 0.0008%, 0.035%, and 0.0009%, respectively. The steady shape of the cell is assumed to be obtained when the difference |**x**(τ+10^5^τ_0_) −**x**(τ)| between the positions of the same node on the membrane at two different moments, separated by 10^5^ timesteps is less than 10^−5^. The calculations are performed on a cluster consisting of 24 Intel@Core i5 processors with 16 GB RAM per node. OpenMP directives are used to parallelize the matrix-vector product computation. Each configuration of the steady shapes is completed via 10^7^ iterations. It is important to note that some of the cases reported in the phase diagrams in results section ran over more than 6 hours on a 12-core node since we decided to avoid using any cutoff or periodic boundary conditions in our system, due to the long-range nature of the hydrodynamic interaction. The use of the appropriate Green’s function (imposing the velocity vanishes on the walls) allowed us to avoid finite size effects, since we can literally consider an infinite domain along the flow direction.

As a validation of our numerical algorithm, the full phase diagram of dynamics of amoeboid swimming have been obtained by both the boundary integral method and the immersed boundary method (Wu *et al.* 2015; 2016). The two methods give the exact same results. In this paper, we explore the ranges of parameters that result in interesting cell shapes, such as the the pear and kidney shapes. In order to have a reference for the conversion of dimensionless units into physical ones, the following dimensional numbers for cell blebs can be used: *R*_0_ = 6*µm*, *µ* = 10^−3^Pa ⋅ s and κ = 10^−18^*J*. This leads to a characteristic time of shape relaxation of about τ_m_ ≈ 0.2*s*, which is consistent with measured values for some cells, such as the amoeboid cells in (Arroyo *et al.* 2012).

### E An accurate algorithm to determine the interior and exterior of a cell

There are two existing algorithms, the ray casting algorithm and the winding number algorithm, to judge whether one target point is located inside or outside a 2D polygon. Here we propose a simpler but equally precise algorithm. The idea is to use the fact that the sum of all the acute angles formed by the target point and every pair of points that define a segment on the two dimensional membrane contour is equal to 2π, if and only if, the target point is inside the cell. Here we assume that acute angles are positive when uniting lines in the clockwise direction. This 2D directional summation of the acute angles formed at the target point is equivalent to calculate the winding number. But our method can be easily generalized to the 3D case, the only change is that the criterion of 2π needs to be changed to 4π, sterad with a summation implemented on all the solid angles formed by the target point and every three border points on the triangulized 2D membrane surface. The winding number algorithm cannot be extended to the 3D case.

## Footnotes

1 This definition of the mean curvature is predicated on choosing the outward normal as the normal to the surface.

2 See Appendix A for a sketch of the derivation of these equations. When the bending and Gaussian moduli are constant the equation for the normal deformation has been derived by Ou-Yang & Helfrich (1989), Capovilla *et al.* (2003), Tu & Ou-Yang (2004) and others.

